# Transposable element insertions are associated with Batesian mimicry in the pantropical butterfly *Hypolimnas misippus*

**DOI:** 10.1101/2023.07.10.548380

**Authors:** Anna Orteu, Marek Kucka, Eunice Katili, Charo Ngumbao, Ian J. Gordon, Ivy Ng’iru, Eva van der Heijden, Gerard Talavera, Ian A. Warren, Steve Collins, Richard H. ffrench-Constant, Dino J. Martins, Yingguang Frank Chan, Chris D. Jiggins, Simon H. Martin

**Affiliations:** Department of Zoology, University of Cambridge, Cambridge, CB2 3EJ, United Kingdom; Tree of Life Programme, Wellcome Sanger Institute, Hinxton, United Kingdom; Friedrich Miescher Laboratory of the Max Planck Society, Tübingen, Germany; Centre of Excellence in Biodiversity, University of Rwanda, Huye, Rwanda; Mpala Research Centre, P.O. Box 555, 10400, Nanyuki, Laikipia, Kenya; School of Biosciences, Cardiff University, Cardiff, CF 10 3AX, UK; UK Centre for Ecology and Hydrology, Wallingford, OX10 8BB, UK; Institut Botànic de Barcelona (IBB, CSIC-Ajuntament de Barcelona), Barcelona, Catalonia, Spain; African Butterfly Research Institute, Nairobi, Kenya; Centre for Ecology and Conservation, University of Exeter in Cornwall, Penryn, TR10 9FE, UK; Turkana Basin Institute, Stony Brook University, Stony Brook, New York, 11794, USA; Institute of Evolutionary Biology, University of Edinburgh, Edinburgh, United Kingdom

**Author notes:** Author for Correspondence: Anna Orteu, Tree of Life Programme, Wellcome Sanger Institute, Hinxton, United Kingdom.

## Abstract

*Hypolimnas misippus* is a Batesian mimic of the toxic African Queen butterfly *(Danaus chrysippus)*. Female *H. misippus* butterflies use two major wing patterning loci (M and A) to imitate the four colour morphs of *D. chrysippus* found in different regions of Africa. In this study, we examine the evolution of the M locus and identify it as an example of adaptive atavism. This phenomenon involves a morphological reversion to an ancestral character that results in an adaptive phenotype. We show that *H. misippus* has re-evolved a wing pattern present in other *Hypolimnas* species for Batesian mimicry of a *D. chrysippus* morph. Using haplotagging, a linked-read sequencing technology, we discover two large transposable element (TE) insertions located at the M locus and establish that these insertions are present in the dominant allele responsible for producing the ancestral and mimetic phenotype. By conducting a comparative analysis involving additional *Hypolimnas* species, we demonstrate that the dominant allele is derived. This suggests that the TEs disrupt a cis-regulatory element, leading to the reversion to an ancestral phenotype that is then utilized for Batesian mimicry of a distinct model, a different morph of *D. chrysippus*. Our findings present a compelling instance of convergent evolution and adaptive atavism, in which the same pattern element has independently evolved multiple times in *Hypolimnas* butterflies, repeatedly playing a role in Batesian mimicry of diverse model species.

## Introduction

Butterfly wing patterns are a classic example of adaptive evolution. Evolutionary genetic studies have dissected the loci controlling wing pattern in several species of butterflies from a wide range of ecotypes and families, providing extensive information on the evolution of adaptive traits (Jiggins 2017; Beldade and Brakefield 2018). The genetic architectures uncovered are varied, from supergenes formed by inversions encompassing multiple loci in *Heliconius numata* (Joron et al. 2011), to transposable element insertions in the peppered moth (van’t Hof et al. 2016) or alternatively spliced isoforms in swallowtails (Kunte et al. 2014).

The *Hypolimnas* genus of tropical butterflies is diverse in wing pattern phenotypes (Figure 1B). Interestingly, the genus presents many instances of Batesian mimicry, with the main models being Danaid species of the *Danaus, Amauris* and *Euploea* genera (Vane-Wright et al. 1977). Despite the diversity in phenotype and model species being mimicked, some wing patten elements are common in most *Hypolimnas*, exemplified by the black-and-white forewing tips found in most species (18/21 species with phenotype data) or the common black or brown background colour. *Hypolimnas* therefore offer an opportunity to study the repeated evolution of adaptive phenotypes in a group that has not been well studied to date.

**Figure 1.**
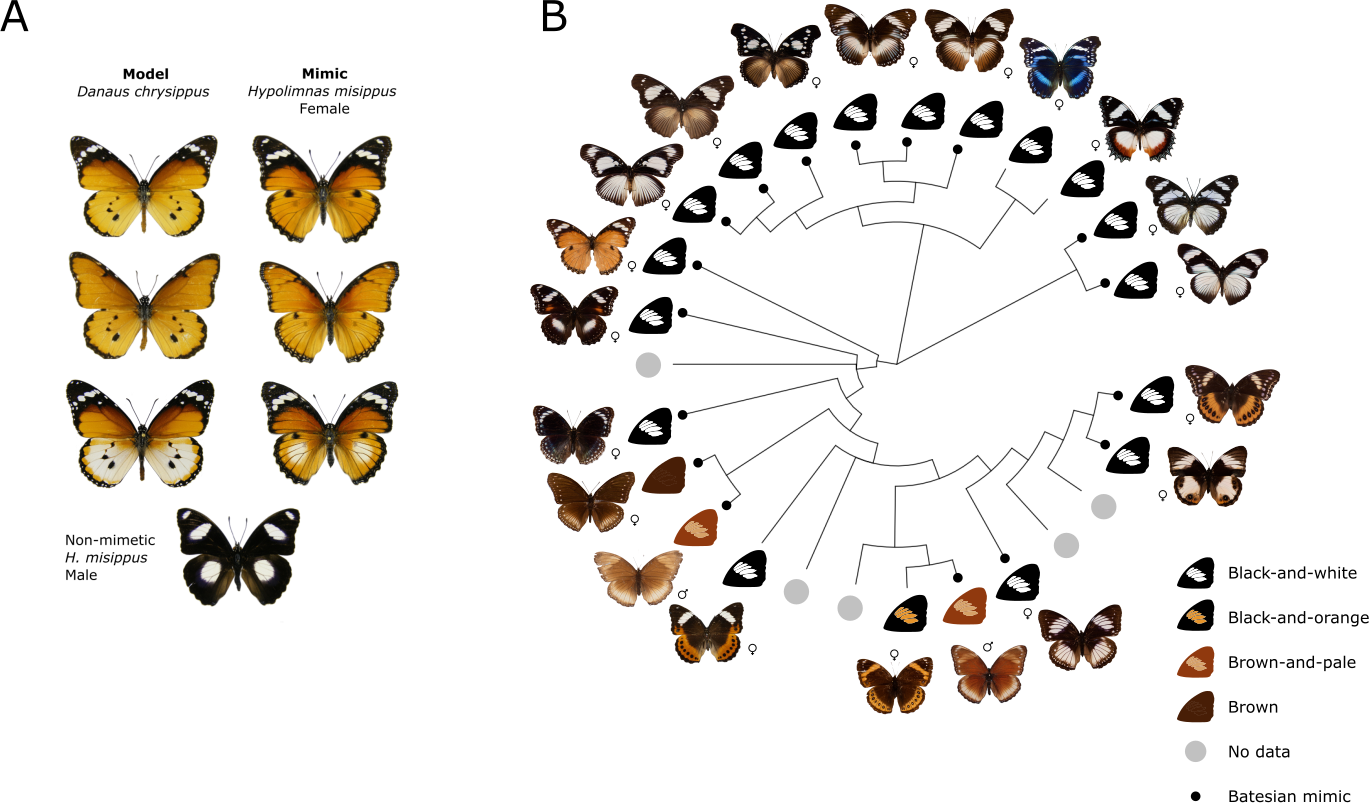
Mimicry in *Hypolimnas missippus* and the *Hypolimnas* genus. **A.** Three female morphs of *H. misippus* side by side with their matching model morphs of *Danaus chrysippus*. Non-mimetic male at the bottom. **B.** Phylogram of the *Hypolimnas* genus. Species showing Batesian mimicry are indicated by a black dot. Recurrent forewing phenotypes indicated by wing drawings. Male and female signs indicate the sex of the individual photographed.

*Hypolimnas misippus* or Diadem is a pantropical butterfly with complex Batesian mimicry. Females are mimetic and polymorphic, with detailed resemblances to four morphs of the toxic African Queen, *Danaus chrysippus* (Figure 1A)(Smith 1973). Despite the striking mimicry, a puzzling mismatch exists in the geographical distribution of *H. misippus* and *D. chrysippus* morphs across Africa, in that the most abundant models are not reflected in the frequency of mimics at a given location (Gordon et al. 2010a). This, together with the fact that maladaptive intermediate morphs of *H. misippus* are commonly found, suggests current selection for mimicry might be weak and raises the question of how the polymorphism is maintained (Gordon and Smith 1998; Gordon et al. 2010a). Clarifying the genetic underpinnings of wing mimicry in *H. misippus* will shed light on this complex case of Batesian mimicry and the forces maintaining polymorphism in the population.

Wing colouration in *H. misippus* is determined by two loci of large effect, the M and A loci, determining forewing and hindwing pattern respectively (Smith and Gordon 1987; Gordon and Smith 1989; VanKuren et al. 2019). The existence of a third locus, the hindwing white suppressor S, has also been hypothesised (Gordon and Smith 1989). The M locus is a Mendelian locus with two alleles, with the dominant M allele (diploid genotype *M-*) producing the mimetic black and white forewing tips in the *misippus* morph; whereas recessive homozygotes (*mm*) have mimetic orange or intermediate forewings, known as the *inaria* and *immima* morphs respectively. Epistasis exists between the M and the A locus, producing the intermediate *immima* forms in *mm* genotypes when the dominant *A* allele for white hindwings is present. Previous work has identified the M locus to an intergenic region of 10 kb near genes of interest such as *pink* and *Sox 5/6* (VanKuren et al. 2019). However, not much is known about the structure of the locus itself, which of the alleles is derived, and whether it arose through de novo mutation or introgression.

Structural variation forms a large part of the genetic variation observed in wild populations and can play a key role in adaptation and speciation (Auton et al. 2015; Wellenreuther et al. 2019). Structural variants (SVs) are typically defined as events larger than 50 bp and include various combinations of gains, losses and rearrangement of genetic material, which can have extensive effects on gene content, as well as genetic contiguity (reviewed in (Ho et al. 2020)). These effects have major roles in adaptation and speciation in many species (reviewed in (Hoffmann and Rieseberg 2008; Kondrashov 2012; Faria et al. 2019)) as well as human disease (Weischenfeldt et al. 2013; Zeevi et al. 2019). For example, inversions have often been associated with complex phenotypes, as reduced recombination at the inversion promotes the joint inheritance of co-adapted alleles (Kirkpatrick and Barton 2006). Examples of this are seen in elytra colouration in ladybirds and reproductive morph switches in the ruff ((Küpper et al. 2015; Lamichhaney et al. 2016; Ando et al. 2018; Gautier et al. 2018); reviewed in (Thompson and Jiggins 2014; Küpper et al. 2015; Orteu and Jiggins 2020). In other cases, gene duplications might give rise to adaptive loci through neo-functionalisation as seen in heterostyly in *Primula* plants and in the complex phenotypes of the wood tiger moth (Li et al. 2016; Brien et al. 2022). Structural variants are therefore crucial in the evolution of adaptive traits and improvements in sequencing techniques are accelerating our ability to detect and study them.

Despite the importance of SVs in phenotypic variation, their study is limited by the difficulty of detecting them using high throughput “short-read” DNA sequencing (Mahmoud et al. 2019). SVs involve the rearrangement of otherwise identical DNA sequences, so their detection often requires sequencing molecules that span the rearranged sequence junction. Relative to the size of an SV (often > 50kb), the fraction of read molecules (typically 300-500 bp) that span junctions can be vanishingly small. This problem is made worse by ambiguous mapping due to repetitive elements, which contribute to the formations of SVs (Sharp et al. 2005; Carvalho and Lupski 2016; Payer et al. 2017). Nevertheless, a number of programs exist to detect SVs from short-read sequencing (Rausch et al. 2012; Sindi et al. 2012; Layer et al. 2014; Iakovishina et al. 2016). Long-read sequencing, in contrast, has improved our power to detect SVs via reads that span repetitive and problematic regions, but is limited by cost (Sedlazeck et al. 2018; Ho et al. 2020).

Linked-read sequencing has emerged as an alternative that combines the scalability of short-read sequencing while retaining linkage information (Marks et al. 2019). The newly developed ‘haplotagging’ is a simple, linked-read technique that can be used to sequence entire study populations with hundreds of individuals (Meier et al. 2021). In this approach, large DNA molecules are barcoded as they are broken up for short-read sequencing. For the purpose of SV detection, the barcoded, larger DNA molecule greatly boosts the fraction of junction-spanning molecules, thus improving detection power. Importantly, happlotagging can be easily scaled up to population level by multiplexing, which makes it possible to track the frequency of polymorphic structural variants in single individuals, making it an ideal tool for the study of adaptation and speciation in non-model organisms (Meier et al. 2021).

Here, we dissect the genetic architecture of an adaptive polymorphism in *H. misippus* using haplotagging data. First, we perform an association study using hundreds of whole genome haplotagging sequences to pinpoint the candidate locus. Then, we use the linked-read information to dissect the genetic structure of the locus by applying Wrath (WRapped Analysis of Tagged Haplotypes, github.com/annaorteu/wrath), our custom program for the analysis of haplotagging data, which validate using published haplotagging data with known SVs. And finally, we perform a cross-species comparison within the genus *Hypolimnas* to investigate the evolutionary history of the adaptive alleles.

## Results

### Forewing mimicry in *Hypolimnas misippus* is controlled by the M locus

To explore the genetic underpinnings of Batesian mimicry in *Hypolimnas misippus*, we first confirmed the previously described identity of the M locus (VanKuren et al. 2019). We sequenced 335 *Hypolimnas misippus* females collected in Kenya and other locations in Africa using haplotagging. The dataset contains 277 individuals and 54 *inaria/immima* (*mm*), sequenced to 0.81 coverage on average (Supplementary Table 1 and Supplementary Figure 1). By using haplotagging with a large dataset, we could sequence to low coverage per individual without compromising statistical power to detect loci associated with mimicry. This is because, although read coverage is low, molecular coverage (i.e. coverage of DNA molecules) is higher in linked read data, as SNP information of reads belonging to the same DNA molecule can be used for imputation and phasing (Marks et al. 2019; Meier et al. 2021). Also, using a large population sample (>200 individuals) facilitates the identification of regions associated with the trait of interest (Lou et al. 2021).

First, we parsed and demultiplexed the data and then imputed SNPs and phased haplotypes, which resulted in the identification of 46.1 M SNPs, with a mean phased block N50 of 109.02 kbp (Supplementary Figure 2). We then performed a GWAS to identify the locus controlling forewing mimicry. A single large peak of association with variation in forewing phenotype was found on chromosome 26 (6,731,000 - 6,743,400 bp; lll^2^ (1, *N=*331)=118; *P-value*=1.703e-27; Figure 2A), corresponding to the *M* locus ((VanKuren et al. 2019), SGBE01001249.1 *Hypolimnas misippus* isolate hmisRef-201410 scaffold88 in UofC_Hmis_v1.0). Principal Component Analysis of the whole of chromosome 26 showed no evidence for population structure in the data (Supplementary Figure 3). In contrast, when using just the associated region for PCA, individuals of the same phenotype were found closer together (Supplementary Figure 4). Closer examination of the associated region in the GWAS result revealed 3 separate peaks of association (Figure 2D), suggestive of a linked haplotype block. By contrast, the intervening SNPs with little to no association occur within tracts with extremely low read depth in individuals with the *inaria* (*mm*) phenotype (Figure 2E), indicative of an absence of sequence reads matching these tracts in *inaria* individuals. One possibility is that structural variation is segregating at this locus.

**Figure 2.**
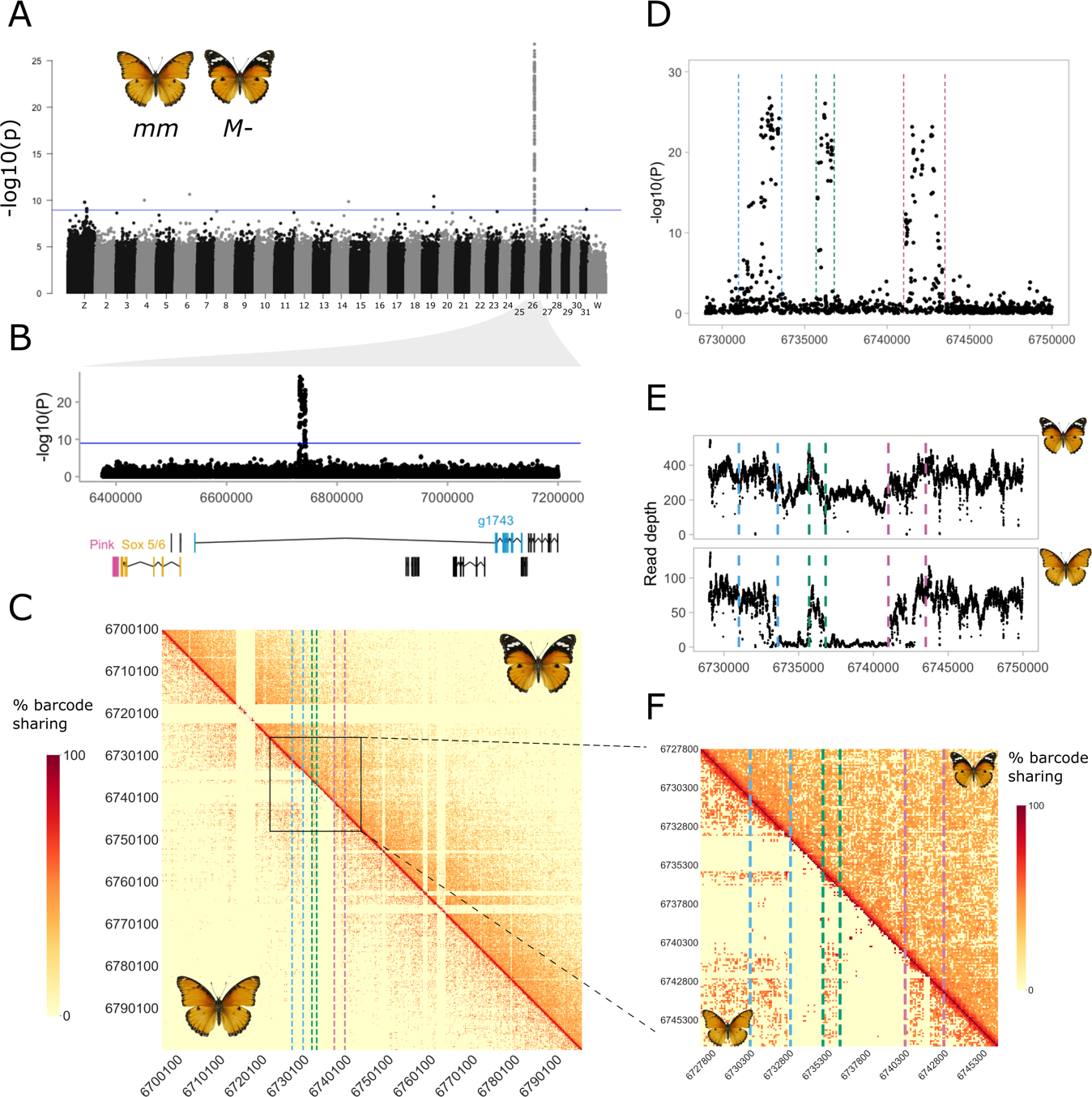
Two large indels are found at the locus associated with forewing mimicry in *mm* individuals. **A.** GWAS of forewing phenotype shows a unique peak at chromosome 26. Blue line indicates the genome wide significance threshold with Bonferroni correction for multiple testing (p=0.05, N=46,088,305) **B.** The GWAS peak showing local annotation track with the three main candidate genes coloured. *Pink* shown in pink, *Sox 5/6* in yellow and *g1743* in blue. **C.** Heatmap of barcode sharing of the region around the association peak. Upper triangle shows barcode sharing for *M-* individuals and lower triangle *mm* individuals. A different pattern of barcode sharing between *mm* and *M-* individuals is seen at the associated region. **D.** At a finer scale the association peak reveals a three-peak structure. **E.** Depth of read coverage around the associated region supports the hypothesis of deletions, as *mm* individuals present almost 0 coverage between the association peaks, while *M-* individuals have more constant coverage throughout the region (note that some *M-* individuals are likely to be heterozygous, carrying one copy of the recessive *m* allele, explaining the partial reduction in read depth in *M-* individuals). **E.** A zoom in of the barcode sharing heatmap reveals a signal of depletion between the peaks of association in *mm* individuals, a signature of deletions relative to the reference (or insertions in the reference).

To explore structural variation more closely, we developed a program for the visualisation of haplotagging data and exploration of structural variants, named Wrath (WRapped Analysis of Tagged Haplotypes). Briefly, Wrath divides chromosomes into genomic windows and quantifies barcode sharing among them, creating a matrix and heatmap that can be used to identify outlier regions caused by structural variants (see Methods).

### *misippus* individuals carry multiple TE insertions at the M locus

The region associated with differences in forewing pattern spans 10 kb in length, thus a very small window size is necessary to elucidate whether there is any structural variation at the locus, given that SVs can only be detected if they are smaller than the genomic windows used (Methods). We visualised barcode sharing using a window size of 100 bp around the *M* locus and identified two putative deletions in the recessive *m* allele relative to the reference genome, which is a haploid assembly generated from an *M* homozygote individual (Figure 2C, F; see Methods). These two indels perfectly match the locations of the troughs of association seen in the GWAS analysis where read coverage is almost zero in *mm* individuals, supporting the hypothesis of two deletions in the *m* allele (or two insertions in the *M* allele) (Figure 2E). This explains the decline in association in these two regions, as SNPs cannot be confidently called in *mm* individuals.

To verify the presence of these indels, we designed PCR primers flanking each indel, and at the breakpoints, and amplified them from *misippus (M-)* and *inaria/immima* (*mm)* individuals (3 individuals per phenotype; Supplementary Figure 5). This confirmed that two insertions of 2.4kb and 4.3kb are present in the dominant *misippus* phenotype relative to the recessive *inaria/immima.* A set of transposable elements (TE) insertions detected with RepeatMasker compose the entirety of the two insertions, which are situated in a 3’-UTR intron of the gene g16415, an ankyrin repeat and sterile alpha motif domain containing gene of unknown function (Figure 2B and 3A and Supplementary Table 2 and 3). Insertion A (most downstream) is composed of a tandem duplication of *Helitron* family transposable element and an unknown TE, while insertion B is composed of 3 *Helitrons,* 4 LINEs and 2 unknown TEs (Figure 3A and Supplementary Table 4). Given that the insertions are found in the dominant allele, the most plausible explanation is that the insertion is modifying the expression of a nearby gene, either *g16415* or others such as *Sox 5/6* and *pink*, by acting as or affecting existing cis-regulatory elements.

**Figure 3.**
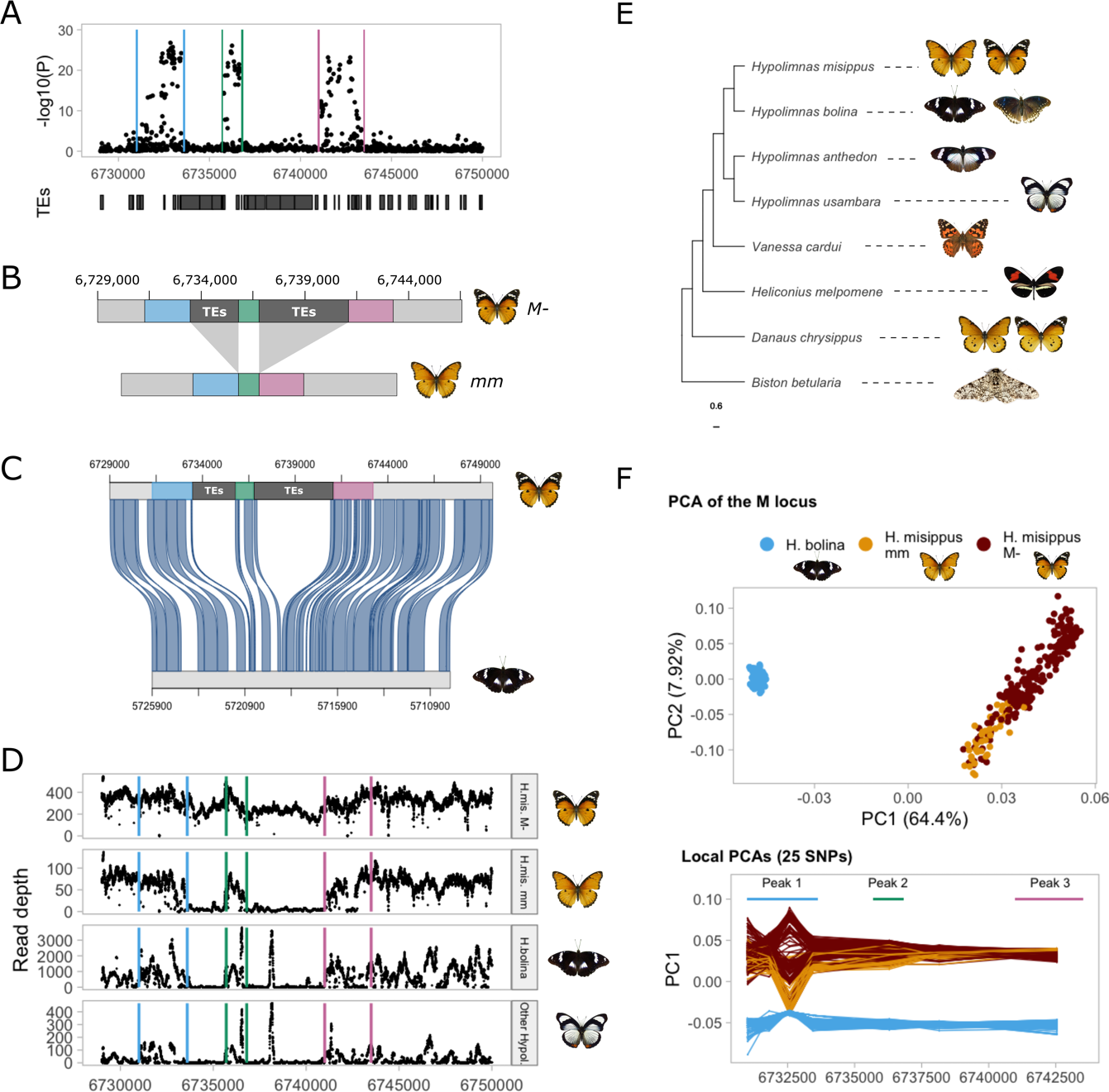
TE insertions are found in the dominant derived allele *m.* **A.** Zoom in of the association peak showing a TE annotation track. The regions between the association peaks are composed by TE insertions. **B.** Schematic of the structure of the dominant derived (top) and recessive ancestral M locus alleles. **C.** The alignment of the *H. misipppus* and *H. bolina* reference genomes shows that *H. bolina* does not present the TE insertions. **D.** Read coverage of *H. bolina* and other *Hypolimnas* species suggests that other *Hypolimnas* do not carry the insertions and that those are thus derived. **E.** A phylogeny of the *Hypolimnas* species used and other Lepidoptera. *H. deceptor* is missing and would be in the same clade as *H. anthedon* and *H. usambara.* Phylogeny extracted from (Kumar et al. 2022). **F.** Principal Component Analysis (PCA) of the locus associated with forewing phenotype reveals structure by species, while local PCA in sliding windows of 25 SNPs reveals that the *m* allele is more similar to the *H. bolina* samples in the region of the first peak (Chromosome 26: 6732649-6732923), supporting the hypothesis of the recessive allele being ancestral.

### The M allele carrying TE insertions is derived and produces an atavistic adaptive phenotype

We next explored the evolutionary history of the M locus. The presence of the TE insertions could either be ancestral or derived. In other words, either the ancestor of both alleles already carried these TE insertions, and they were subsequently deleted to form the *m* allele as it is today, or the ancestor lacked the TE insertions, and they were inserted to form the *M* allele as it is today. To test these hypotheses we explored this region in the genome of *Hypolimnas bolina*, a relative of *H. misippus* with very different wing patterns that diverged approximately 8 million years ago (Sahoo et al. 2018). First, we aligned the *H. misippus* and *H. bolina* reference genomes using Satsuma2, an aligner intended for inferring homology from sequence similarity (Grabherr et al. 2010). We identified an orthologous region on chromosome 26 of *H. bolina* showing strong synteny across the M locus, with no further rearrangements except for the TE insertions, indicating conserved synteny in *H. misippus* (Figure 3B). Furthermore, the alignment shows that, while the peaks of association have homologous sequences in chromosome 26 in *H. bolina,* the two indels between the peaks have no matches in the *H. bolina* genome (Figure 3B). This strongly suggests that the *m* allele is ancestral, and the TEs represent derived insertions into the *M* allele.

To further explore the origin of the alleles, we analysed whole genome resequencing data from 4 other *Hypolimnas* species, including 214 *H. bolina,* 4 *H. anthedon,* 4 *H. deceptor* and 2 *H. usambara* (Figure 3E), sequenced to an average coverage of ∼6.5X. First, we mapped all resequenced *Hypolimnas* to the *H. misippus* reference genome and quantified read coverage at the M locus. Read coverage across the TE insertions is approximately zero in the outgroup species (Figure 3D), confirming the hypothesis that the TE insertions are derived and unique to the *M* allele of *H. missippus*.

To investigate in more detail the evolution of the *M* and *m* alleles, we evaluated the SNP variation at the locus across all the *H. misippus* and *H. bolina* samples using Principal Component Analysis (PCA). PCA was conducted using the SNPs of the associated region, and then repeated using windows of 25 SNPs across the region. We chose to use PCA because compared to other techniques such as phylogenetic trees, PCA or similar dimension reduction techniques focus on the main mode of variation and can be more robust under low-coverage sequencing scenarios such as our dataset. Here, our aim is to identify overall genetic similarity: in broader regions, a PCA should recapitulate the species relationships and separate all *H. misippus* from all *H. bolina* samples. However, if the derived M allele accumulated an excess of derived mutations, we may find certain tracts in which the *inaria/immima* individuals, which carry the ancestral *m*-allele, group more closely with the *H. bolina* individuals than with the *H. missipus* individuals carrying the *M allele*.

Across the whole associated region that contains the M locus, the PCA reflects the species relationships, with one cluster for each species (Figure 3F top). Contrastingly, performing local PCAs in genomic windows of 25 SNPs reveals a different pattern. We identified a region containing the top associated SNPs of the most upstream peak of association (6732649-6732923) for which the relationship between the samples did not reflect the species relationship (Figure 3F bottom). *Inaria/immima mm* individuals are found closer to *H. bolina* than to *misippus M-* individuals, which could suggest that the *m* allele is ancestral. The TE insertions found in the *M* allele have reduced recombination, because of reduced effective population size, which could lead to the reduction of recombination at the flanking regions. The TEs might be disrupting a functional element and thus under weak selection which could be coupled with the low recombination and lead to the accumulation of mutations. These coupled effects could make the *M* allele retain fewer ancestral SNPs at the flanking region than the *m* allele. Taken all together, the read-coverage, reference genome alignment and local PCA at the associated region suggest that the recessive m allele that produces orange forewings in homozygosis is ancestral to the dominant *M* allele that produces black-and-white forewing tips.

### Visualisation of inversions, deletions, translocations, and duplications in a dataset of wild *Heliconius* butterflies using Wrath

We further tested the performance of Wrath, our program for the visualisation and exploration of SVs from haplotagging data, in a different system, applying it to an existing haplotagging dataset of the two tropical butterfly species, *Heliconius erato* and *melpomene* (Meier et al. 2021). Each species presents two morphs or subspecies with mimetic wing patterns which hybridise: *H. melpomene plesseni* and *malleti* and *H. erato notabilis* and *lativitta*. We used this dataset to explore which SVs are found in these populations. First, we searched for any SVs present in the dataset genome-wide and identified 3,072 large (>50 kb) putative SVs in *H. melpomene* and 2,885 in *H. erato*. We then explored these using the heatmaps produced by Wrath (Figure 4). Patterns of barcode sharing observed in the heatmaps can be used to identify the type of SV present in the samples. For example, inversions result in a bowtie pattern in the heatmap, as more barcodes are shared than expected between loci that are far apart in the reference genome (Figure 4). We produced heatmap plots for all large scaffolds of *H. melpomene* and *H. erato* and explored the SVs present in the dataset (Figure 4C, Supplementary Data). For example, a known SV in chromosome 2 in *H. erato* was clearly visible in the heatmap (Supplementary Figure 6). With this, we show that Wrath can visualise patterns of barcode sharing and help prioritise the order of exploration of SVs, as visual examination of the haplotagging data helps explore the SV content in the samples. Nonetheless, some SVs produce similar or matching signals. For example, patterns of interchromosomal translocations like that shown in Figure 4A can also be produced by TE insertions.

**Figure 4.**
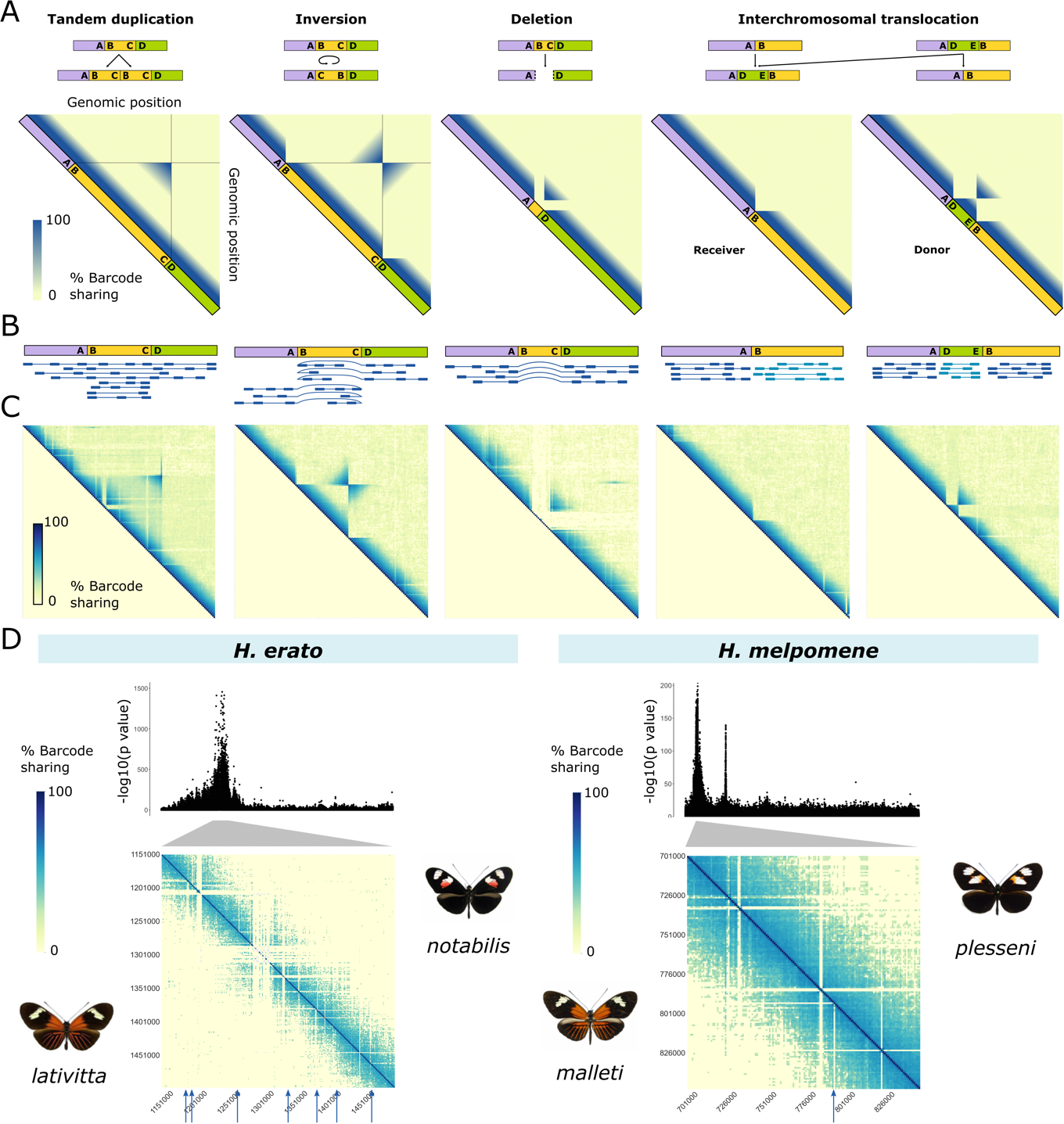
Wrath produces visualisations of haplotagging data to identify candidate structural variants. **A.** Hypothetical Wrath outputs for some SV types showing heatmaps of barcode sharing between genomic windows of a chromosome. On top of each heatmap are depicted the reference genome and its order of loci and below a depiction of the rearranged genome region containing an SV. Points A, B, C and D depict different loci around the breakpoints of the SV that we use as a guide through the diagrams. **B.** Linked-read mapping pattern on the reference genome for each of the hypothetical SVs. **C.** Wrath output heatmaps from the *Heliconius* dataset depicting possible SVs that match the hypothetical predictions. **D.** Wrath outputs of the region around one of the loci (the *optix* locus) associated with colour pattern in *H. erato* (left) and *H. melpomene* (right). Each triangle half of the matrix depicts barcode sharing for one of the colour pattern subspecies—in each case depicted by the side. Blue arrows point at putative deletions that are polymorphic between the two subspecies. Above the heatmap is plotted the Manhattan plot of the GWAS association of colour pattern between each pair of subspecies. These show only the region around the associated *optix* locus. Grey triangles depict the correspondence of regions between the GWAS and heatmap.

Finally, we assessed the time required to run on the dataset. In a small subset of the data and running with 20 threads, the parallel implementation ran 10x faster than a single-threaded implementation of the same algorithm (Supplementary Figure 7).

### Multiple deletions are found at the colour associated locus near *optix* in *H. melpomene* and *H. erato*

We were interested to know whether known wing patterning loci in *Heliconius* were also associated with structural variation. Using Wrath, we found that in both species pairs there are multiple putative deletions of 1-10 kb that differentiate colour morphs at the locus associated with red pattern elements near the gene *optix* (Figure 4D). Deletions leave an area depleted of barcode sharing, as reads do not map to the area, which can be visually identified using the heatmaps (Figure 4A-C). However, only polymorphic deletions can be identified, as deletions that are fixed in all sequenced individuals but not in the reference cannot be distinguished from assembly artefacts or poor mapping (e.g., repetitive regions for which mapping reads are filtered out due to low mapping quality). Larger deletions (>10 kb) leave an additional signature in the heatmap, showing increased barcode sharing between the breakpoints, in the shape of a triangle (Figure 4A-C). This is because more molecules than expected span through the breakpoints in individuals presenting the deletion. This signature is harder to detect in smaller deletions, as barcode sharing across the small, deleted region is similarly high in individuals with and without the deletion.

The red patterning locus contains cis-regulatory elements that control the expression of *optix* and influence development of red colour elements, which have been functionally tested with CRISPR (Lewis et al. 2019). Thus, one possibility is that these deletions are disrupting the function of an *optix* CRE and affecting *optix* expression, although they will need to be functionally tested. Alternatively, it could be that these deletions are in linkage with selected SNPs and thus associated with the adaptive colour patterns. Finally, the deletion detected in *H. melpomene* was detected previously using short-read sequencing of two different sub-species (Wallbank et al. 2016).

## Discussion

Here, we present a case of adaptive atavism in the diadem butterfly, *Hypolimnas misippus*, in which the derived allele is associated with a reversionreversion to an ancestral yet adaptive phenotype. Atavisms are caused by mutational or recombination events that enable the pre-existing developmental machinery to reproduce the ancestral character (REF). Crucially, they are often maladaptive, as the lost phenotype has been selected against, such as hind limbs in whales and teeth in birds, or are associated with a malfunctional state such as cancer (Thomas et al. 2017). In line with this, Stephen Jay Gould coined the term Dollo’s law to refer to the paleontological observation that morphological traits that are lost in an evolutionary lineage, do not later on re-evolve in that lineage (Gould 1970). We present a case where the atavistic phenotype is adaptive, with the derived allele of the M locus in *H. misippus* producing a mimetic wing phenotype. We show that two large insertions of 2.4kb and 4.3kb are found in the dominant allele of the M locus and that these are formed by multiple TE insertions. By comparison to other *Hypolimnas* species, we show that the insertions are derived, suggesting that the orange wing phenotype evolved first and that insertions at the M locus subsequently reverted the phenotype to a more *Nymphalidae*-like phenotype. Melanised apexes in the forewing with subapical white bands are a common wing phenotype in *Hypolimnas* present in 81% of the species (17 out of 21 with phenotype data; Figure 1B) and in Nymphalids such as Danaids or some *Nymphalinae,* including *Antanartia* and *Vanessa species* (e.g., *Vanessa cardui,* Figure 4.3D). Here, we can understand the phenomenon of “evolutionary reversion” in the *H. misippus* butterfly as a molecular example of convergent evolution, that is the re-evolution of the same phenotype via regulatory rewiring. We present an example of an adaptive atavistic phenotype in which the derived M allele is producing an adaptive atavistic phenotype that causes a reversal to an ancestral phenotype, the black-and-white phenotype, with an adaptive function in Batesian mimicry. Adaptive atavism is a rare event with only a few known examples such as the re-evolution of wings in stick insects (Whiting et al. 2003) and aphids (Saleh Ziabari et al. 2023), sexual reproduction in oribatid mites (Domes et al. 2007), and shell coiling in gastropods (Collin and Cipriani 2003).

Alternatively, it could be that the TE insertions are not directly causal but in linkage disequilibrium with the causal mutation. Functional testing such as gene editing with CRISPR-Cas9 would be necessary to prove the causality of the TE insertions. Furthermore, introgression of the *m* allele from another Hypolimnas species such as *H. bolina* could explain the local PCA results. However, phylogenetic trees of those SNPs do not show any introgression signal, that is clustering of *H. bolina* samples with *H. misippus m* alleles (Supplementary Figure 9).

The TE insertions identified are found in the intron of the gene g16415, which encodes an ankyrin repeat and sterile alpha motif domain containing protein of unknown function. Our results represent a similar case to the peppered moth, where a 10-kb TE insertion increases the expression of *cortex*, resulting in the production of melanic morphs (van’t Hof et al. 2016). Similarly, variation in wing colour in the *Heliconius melpomene/timareta* lineage is associated with a TE insertion in the *cis*-regulatory region of *cortex,* suggesting that cis-regulatory structural variation controls these mimetic phenotypes (Livraghi et al. 2021). Outside of Lepidoptera, TE insertions in cis-regulatory regions have also been found to be of adaptive importance, such as in egg-spot phenotypes in cichlid fish (Santos et al. 2014) and flowering time in the an annual and inbreeding forb *Capsella rubella* (Niu et al. 2019). Given that the M locus insertion is found in a non-coding region, the most plausible explanation is that the insertion is modifying the expression of a nearby gene, either g16415 or others such as *Sox 5/6* and *pink* by acting as or affecting existing cis-regulatory elements. Also, given that the insertion is found in the dominant allele, an increase in expression of the candidate gene or a certain isoform are possible explanations, which could be achieved by the disruption of a repressor or the generation of a novel enhancing function. Overall, this case adds more evidence that cis-regulatory mutations are associated with pattern variation, while coding mutations are more likely to be associated with colour, and sheds light on the adaptive importance of TEs (Casacuberta and González 2013; van’t Hof et al. 2016; Orteu and Jiggins 2020).

Although the accuracy of the mimicry seen in *H. misippus* would suggest strong selection, intermediates are often found (Gordon et al. 2010a). Moreover, while the subspecies of the model *D. chrysippus* each have large geographically distinct regions monomorphism (Liu et al. 2022), all four of the *H. misippus* mimetic morphs are found in each of these regions. In other words, there is a phenotypic mismatch, in which many *H. misippus* individuals are mimics of a model that is rare or absent in their region, suggesting that other evolutionary forces might be at play (Gordon et al. 2010a). Negative-frequency dependent selection has been invoked as one of several forces maintaining the polymorphism in *H. misippus* (Gordon 1987; Gordon et al. 2010a). Another alternative is that structural variation causes associative overdominance and prevents the fixation of a single allele even in scenarios of weak mimicry selection. An example of this has been shown in *Heliconius numata* in which a supergene containing three inversions controls wing phenotype (Jay et al. 2021). The inversions result in a region of low recombination, which in turn lead to the accumulation of deleterious recessive mutations. This leads to reduced fitness of homozygotes for both inversion alleles due to the deleterious effects of recessive mutations at different, but tightly linked sites (i.e., associative overdominance). The TE insertions at the M of *H. misippus* could theoretically lead to associative overdominance by reducing local recombination between M and m alleles, but the impact on recombination is likely to be less severe than that of a large inversion, so the role of recombination suppression on the maintenance of polymorphism requires further investigation.

In addition to our empirical results, we also present Wrath, a user-friendly, flexible, and fast tool for the visualisation of haplotagging data and exploration candidate SVs, which we test using two large haplotagging datasets. Wrath produces heatmap plots of barcode sharing that can be used to visually inspect the data and identify candidate SVs. Wrath can be run with any chosen window size, which gives flexibility to the user and allows for the detection of SVs of different sizes. There are multiple software solutions for the identification of SVs from linked-read data, including LongRanger (Sudmant et al. 2015), Leviathan (Morisse et al. 2021), NAIBR (Elyanow et al. 2018) and GROC-SV (Spies et al. 2017). LongRanger and GROC-SV are very well curated tools for the analysis of linked-reads produced by 10X-Genomics, while NAIBR uses the BAM files produced by LongRanger for its SV detection pipeline. Whilst these programs could be used for haplotagging data, the data would need to be converted to the 10X-Genomics format for input to these programs. Finally, Leviathan can take haplotagging data as input and produces a list of detected candidate SVs and their predicted breakpoints, however, unlike the above tools, Leviathan does not produce graphic visualisations of barcode sharing, which we have found a very useful tool for manual verification of putative SVs. We tested the effectiveness of Wrath using our dataset of *Hypolimnas misippus* and a published dataset from *Heliconius* butterflies and show that Wrath can be useful in the visualisation of haplotagging data to identify candidate SVs.

Altogether, our study presents a striking case of adaptive atavism in which an ancestral trait reappears for an adaptive function. Furthermore, our results highlight the importance of structural variation in the evolution of adaptive phenotypes, adding to the mounting evidence that TEs have an important role in adaptive evolution and particularly in the evolution of colour and mimetic phenotypes in Lepidoptera. Finally, we have shown that Wrath is an easy and flexible means to visualise haplotagging data and explore candidate SVs.

## Methods

### Visualisation and exploration of SVs from haplotagging data using Wrath

To analyse haplotagging data, we developed Wrath (WRapped Analysis of Tagged Haplotypes, available at github.com/annaorteu/wrath), a program for the exploration and visualisation of SVs consisting of three steps.

### Barcode parsing

Haplotagging reads are produced using magnetic beads that present a modified Tn5 enzyme on their surface carrying sequencing adapters, each with a unique barcode. During library preparation, DNA molecules wrap around the beads and are cut into smaller fragments and barcodes attached to them. Thus, reads belonging to the same DNA molecule present the same unique barcode, and the small size of the beads ensures that each barcode combination is unique to one or a small number of molecules. In the sequencing files, barcode information is included as four nucleotide sequences of 6 bp each (two per index read). To analyse haplotagging data, first, molecule information needs to be included as a BX tag in the information fields of the fastq files, a process known as molecule demultiplexing. Once the reads include information on their molecule-of-origin in their BX tag, they are ready to be mapped.

### SV visualisation

Using mapped reads and a reference genome, Wrath plots heatmaps of barcode sharing with a single command. Wrath can also produce a list of candidate SVs. Haplotagging reads belonging to the same DNA molecule present the same unique barcode and are expected to map in close proximity in the genome in the absence of rearrangements. Thus, we can use patterns of barcode sharing between more distant genomic windows that exceed the background expectation to identify SVs.

Wrath divides a given chromosome into *n* windows of size *m* (m needs to be specified, by default 10 kb) and identifies the barcodes attached to the reads mapping in each of the genomic windows. Window size is chosen based on two factors. First, computational overhead, as Wrath builds a matrix of *nxn* dimensions which can require a large amount of memory for large values of *n*. And second, molecule size, which depends on several factors such as sample preservation, DNA extraction and library preparation. By default, molecules are assumed to be centred around 50kb in length, although they can be much larger (Meier et al. 2021). Window size needs to be smaller than molecule size (e.g., 10kpb window size and 50 kb molecule size) as the identification of SVs is only possible if molecules span more than one window.

Once the chromosome has been split into windows, Wrath determines the barcodes that are present in each of those windows and calculates the Jaccard index for each pair of windows along the chromosome and stores the value in a matrix of *nxn* dimensions. The Jaccard index is an index of similarity that quantifies barcode sharing between windows:

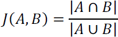

Where J is the Jaccard value between window A and B from a given chromosome.

The highest values are expected around the diagonal, which then decay exponentially with distance from it. This is because windows that are closer to each other are expected to share more barcodes than windows that are further apart, as DNA molecules span more than one window. Structural variants such as moderate to large inversions (>50 kb), intrachromosomal translocations and long duplicated regions are expected to deviate from the background distribution of barcode sharing. For example, inversions show up as bowtie patterns of excessive barcode sharing between windows that are far apart (FIGURE 1). We define excessive sharing as barcode sharing that is statistically higher than that expected by the distance between the windows given that barcode sharing decays exponentially from the diagonal. Conversely, SV of a much smaller size than the average molecule length cannot be detected with linked-reads. In those cases, short-read methods are more appropriate.

Finally, the construction of the matrix and calculation of jaccard indices for each genomic window can be a computationally expensive task, for that Wrath can be run in parallel, which minimizes computational time.

### Exploration of candidate SVs

Wrath has the additional functionality of producing a list of candidate SVs. Wrath detects SVs where there is an excess of barcode sharing, such as in inversions, interchromosomal translocations and duplications. The distance of each entry of the matrix to the diagonal is calculated and a double exponential decay model fitted to the data such that:

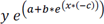

Where *x* is the distance of each entry to the diagonal, *y* is the value of the entry (barcode sharing between a pair of windows, and *a*, *b* and *c* are parameters of the function calculated from the data.

The prediction bands fitted by the model includes the background distribution of barcode sharing and any windows whose values are outside the prediction bands (lll=0.05) are then classified as putative SVs. Once the model has been fitted, Wrath outputs a list of putative SVs with their genomic coordinates and produces plots of the fitted model and identified outliers. The putative SVs are not classified into SV types and are intended to be used for prioritisation processes before further exploration.

Additionally, Wrath scales barcode sharing values according to their distance to the diagonal using Z-scores and filters out any absolute Z-score larger than a given threshold (default 10). This additional filter reduces the number of false positives close to the diagonal, as the closer the genomic windows are, the more variation there is in their barcode sharing.

Wrath can be applied to single populations to visualise and explore putative SVs in each chromosome of the genome. It can also be applied to detect SVs in different populations separately, which can then easily be compared and scanned for overlaps using bedtools (see case-studies below).

### Analyses of *Heliconius* data

Data from (Meier et al. 2021), was used. First, reads were pre-processed, adapters and low-quality ends were trimmed using TRIMMOMATIC (Bolger et al. 2014). Then, we mapped the reads to the respective reference genome, *H. erato* v1.0 (Nadeau et al. 2014) and *H. melpomene* v2.5 (Davey et al. 2016; Davey et al. 2017), using BWA-mem (Li 2013) and marked PCR duplicates using the MarkDuplicates utility from Picard tools (broadinstitute.github.io/picard). Alignment (BAM) files were used as input for Wrath, which we ran separately for each scaffold in the reference genome with a window size of 10 kb. Reads with a mapping quality below 10 were filtered out by Wrath. To examine the colour loci specifically, we ran Wrath with a window size of 1 kb and specifying the desired coordinates.

### Hypolimnas misippus sampling

Samples were collected in multiple locations in Africa. Most sample bodies were preserved in 100% ethanol immediately after collection, with some exceptions that were air dried. Samples in ethanol were kept at room temperature for 2 months and then stored at −80C. A small (1/8) piece of the thorax was used for sequencing.

### *H. misippus* sequencing library construction

Haplotag libraries were prepared essentially as described in Meier et al., 2021, with the following modifications. Broadly, haplotag libraries were prepared in batches of 96 samples. Briefly, genomic DNA sample was diluted to 0.15 ng/µl with 10 mM Tris, pH8 and quantified with Quant-iT PicoGreen dsDNA Assay Kit (Thermo Fisher Scientific). We used only 1.2 µl of haplotagging beads (∼0.88 million beads, each carrying one of 885K well-specific barcodes) per sample; 30 µl of WASH buffer (20 mM Tris pH8, 50 mM NaCl, 0.1% Triton X-100); 10 µl of 5x tagmentation buffer (50 mM TAPS pH 8.5 with NaOH, 25 mM MgCl2, 50% N,N-dimethylformamide); and 25 µl of 0.6% SDS for Tn5-stripping following tagmentation. For sub-sampling, 1/10th of the beads+DNA (0.15 ng DNA per sample) from each of the 96 samples was pooled into a single 8-tube-PCR-strip, and then again from every 8 pools into 4 final samples pools. With only 4 pooled samples on the magnetic stand, the buffer was removed, and 20 µl of 1x Lambda Exonuclease buffer, supplemented with 10 units of Exonuclease we and 5 units of Lambda Exonuclease (New England BioLabs), was added to each sample. Samples were incubated at 37 °C for 30 minutes, and then washed twice for 5 minutes with 150 µl of WASH buffer. DNA library was then amplified using Q5 High-Fidelity DNA Polymerase (New England BioLabs) in four 25 µl PCR reaction according to manufacturer’s instructions, using 4 µl of 10 µM TruSeq-F AATGATACGGCGACCACCGAGATCTACAC and TruSeq-R CAAGCAGAAGACGGCATACGAGAT primers, with the following cycling conditions: 10 min at 72°C followed by 30 sec 98°C and 10 cycles of: 98°C for 15 sec, 65°C for 30 sec and 72°C for 60 sec. Libraries were pooled after PCR into a single library pool, size selected using Ampure magnetic beads (Beckman Coulter), Qubit quantified, and adjusted with 10 mM Tris, pH8, 0.1 mM EDTA to 2.5 nM concentration for sequencing.

Sequencing and demultiplexing. Pooled libraries were sequenced by a HiSeq 3000 (Illumina) instrument at the Genome Core Facility at the MPI Tübingen Campus with a 150+13+12+150 cycle run setting, such that the run produced 13 and 12nt in the i7 and i5 index reads, respectively. Sequence data were first converted into fastq format using bcl2fastq v2.17.1.14 with the following parameters --use-bases-mask=Y150,I13,I12,Y150 --minimum-trimmed-read-length=1 --mask-short-adapter-reads=1 --create-fastq-for-index-reads (Illumina).

### SNP calling and imputation

First, molecules were de-multiplexed. When using haplotagging, the molecule of origin information is embedded in the read name in the fastq file as a string of four barcodes of six nucleotides each. This was done as described in described in Meier et al., 2021 to generate the modified fastq files. The molecule ID is then included in the BX tag of each read. Barcode mismatches caused by sequencing errors are allowed as long as there is an unambiguous closest match. Once the BX tag was created, we pre-processed the reads, cutting adapters and low-quality ends using TRIMMOMATIC, and marking PCR duplicates using the MarkDuplicates utility from Picard tools (broadinstitute.github.io/picard) with two specific options CREATE_INDEX=TRUE and READ_ONE_BARCODE_TAG=BX. We then de-multiplexed the individuals using their barcodes and included their individual ID information in the read group field.

SNPs were identified using the mpileup utility of bcftools v1.11 (Danecek et al. 2021), running each chromosome separately including the INFO/AD,AD,DP,DV,DPR,INFO/DPR,DP4,SP tags in the output (-a option), setting the minimum mapping quality to 10 (-q) and the minimum base quality to 20 (-Q), ignoring Read Group tags (--ignore-RG) and removing duplicates (-F 1024), and the optput directly piped to bcftools call using the alternative model for multiallelic and rare-variant calling (--multiallelic-caller), including only variants in the output (--variants-only) and the fields GQ and GP (-f GQ,GP). Then, using bcftools query (-f), we selected the generated a file containing the chromosome, position, reference and alternative alleles for each SNP and with that produced a file of SNP positions that we could use as one of the inputs for the SNP imputation program STITCH (Davies et al. 2016). Following that we generated genomic windows of 500 kb using bedtools over which we could iterate to run the remainder of the pipeline.

We ran STITCH separately for each of the genomic intervals using all our bam files as input. STITCH imputes SNPs from read and linked-read information but requires fine tuning of the input parameters. To optimise the values, we tested multiple values and compared the results, evaluating their performance using the M locus (Supplementary Methods). SNPs at the M locus are expected to be 0/0 or 0/1 for *misippus* individuals and 1/1 for *inaria* individuals. Options that optimised the results were K=30, method=diploid, nGen=500, readAware=TRUE, keepInterimFiles=FALSE, shuffle_bin_radius=500, expRate=5, iSizeUpperLimit=500000, keepSampleReadsInRAM=TRUE, use_bx_tag=FALSE. We concatenated the resulting imputed variant calls (vcf files) using bcftools concat.

### SNP phasing in *H. misippus*

To phase the SNPs into haplotypes, we used HapCut2 (Edge et al. 2017), which we ran separately for each of the 500 kb intervals used for imputation and each individual separately. First, we filtered SNPs based on their informativeness (INFO_SCORE >= 0.2) and selected all heterozygous SNPs. We used this as input for the --extractHAIRS utility of HapCut2 together with the BAM files with marked duplicates and the option --10X turned on, which indicates that the input contains linked reads. This produced a filed with unliked fragments, which we then used as input for the LinkFragments.py script of HapCut2, which integrates the information of the linked reads. We specified a maximum distance of 50kb. Then, we used the linked fragment file and vcf as input for the HAPCUT2 utility with the option --nf 1 -- threshold 30 --error_analysis_mode 1 --call_homozygous 1. Finally, we integrated the resulting vcf to out vcf of homozygous sites.

### Phenotyping and GWAS

We photographed forewings and hindwings of each individual in a standardised set-up, using a green background and a colour checker. Phenotypes were scored by hand following the phenotype categorisations of (Gordon et al. 2010b), coding *misippus* morphs as 1 and *inaria* as 0. All sample phenotypes are found in Supplementary Table 5.4 (REF). Using the merged HAPCUT2 output vcf file as input and the phenotype scores, we performed a GWAS with Plink v1.9 (Purcell et al. 2007) using the option –assoc.

### Detection of SVs in *H. misippus*

Genome wide SVs were identified using Wrath using the same method as for the *Heliconius* data. We used the intersect utility from bedtools v2.30.0 to assess overlap between SVs identified in homozygous recessive, and heterozygous and homozygous dominant individuals, setting the minimum fraction of overlap to 0.8 for both sets and extracting only one match per SV (intersectBed with options -f 0.8 -F 0.8 -u).

### DNA extractions and amplification of the M locus in *H. misippus*

DNA extractions were carried out using a custom protocol using PureLink buffers and homemade magnetic beads. Briefly, a small piece of thorax tissue (1/10) is placed in a 8-tube PCR trip. Then, 45 uL of PureLink Digestion buffer and 10 uL of Proteinase K (20mg/mL) are added, and the mix is incubated at 58°C with shaking (500 rpm). Thereafter, we added 2uL of RNAseA (DNAse free, 10mg/mL) and incubated it 10min at room temperature. Then, we added 45uL of PureLink Lysis buffer and incubated at 58°C for 30 minutes with shaking (500 rpm). We then used a homemade magnetic bead mix to extract the DNA from the lysate. First, we added 37.5 uL of magnetic beads and 75 uL of lysate to a 96-well plate. After mixing, we incubated 15 minutes at room temperature, placed the plate in a magnetic stand for 10 minutes, removed the supernatant and cleaned the beads with 80% ethanol. After drying out, we added 50uL of 10mM Tris (pH=8) to elute and incubated at 45°C for 15min without resuspending. Then, we resuspended the beads and incubated for 20 minutes at room temperature. Finally, we placed the plate on the magnetic stand and, after 10 minutes, transferred the supernatant (the DNA) to a fresh tube.

To amplify the regions of interest, we designed primers at each side of the deletions and at the breakpoints (SUP FIGURE XX). We used a Q5 High-Fidelity 2X Master Mix from New England BioLabs and with 35 cycles. We used 8 individuals, 3 *inaria/immima* (CAM035230, CAM035232, CAM035239, CAM035240, CAM035244, CAM035245, CAM035249, CAM035250).

### Reference genome alignments

To identify putative homologous regions of the reference genomes of *H. bolina* and *H misippus*, we aligned the two references to each using Satsuma2 (Grabherr et al. 2010) with default parameters. We visualised the resulting alignments using the asynt R functions (Kim et al. 2022).

### Sample preparation and genome wide analysis of *H. bolina* samples and other *Hypolimnas* species

214 wild and reared *H. bolina* samples were used. Briefly, samples DNA was extracted, and DNA Nextera libraries prepared using custom protocols. First, DNA was extracted of the *H. bolina* samples following a custom protocol that uses PureLink buffers and homemade magnetic beads. Briefly, a small piece of thorax tissue was placed in PureLink Digestion buffer and Proteinase K and incubated for 2-3 hours at 58°C with shaking (500rpm). RNAse (DNAse free) was added together with PureLink Lysis Buffer and the samples incubated for 30 min at 58°C with shaking (500 rpm). Afterwards, to pellet any undigested solids, the samples were span at 4000g for 10 at room temperature. Following that, the DNA was extracted from the lysate using a homemade magnetic bead mix with two rounds of 80% ethanol clean-ups.

From the extracted DNA, libraries were prepared following a method based on Nextera DNA Library Prep (Illumina, Inc.) with purified Tn5 transposase (Picelli et al. 2014). PCR extension with the N701–N800 i7-index primer and the N501-N508 and N5017 i5-index primers was performed to barcode the samples. Library purification and size selection was done using the same homemade beads as above.

Short-read data from whole genomes were sequenced to ∼6.5X in coverage. Reads were trimmed using fastp (Chen et al. 2018) and mapped to the reference genome *HypMisi_v2* and *HypMisi_v1* using BWA-MEM2 (Vasimuddin et al. 2019). PCR duplicated were marked using MarkDuplicatesSpark from GATK (Auwera and O’Connor 2020) and SNPs called bcftools v1.11 (Danecek et al. 2021) mpileup like described above. Imputation was carried out with STITCH (Davies et al. 2016) using the same settings as for *H. misippus*.

### Read depth analysis

We calculated read depth from the bam files using the depth utility from samtools (Danecek et al. 2021) with the --a option to output depth for all sites, including those with no reads mapping to them. We visualised the output in R v 4.1.2 using the ggplot2 package (Wickham 2016).

### Principal Component Analysis

VCFs of phased haplotypes were used for PCA, which was performed using Plink v1.9.

## Notes

https://github.com/annaorteu/wrath

### Competing Interest Statement

The authors have declared no competing interest.

